# Frontal theta and posterior alpha oscillations reflect the reactivation of working memory representations following interruptions

**DOI:** 10.1101/2020.08.20.259473

**Authors:** Bianca Zickerick, Marlene Rösner, Melinda Sabo, Daniel Schneider

**Author notes:** Authors contributed equally (Bianca Zickerick, Marlene Rösner). Corresponding author, M.A. Bianca Zickerick, Leibniz Research Centre for Working Environment and Human Factors, Ardeystraße 67, 44139 Dortmund, Germany.

## Abstract

Interruptions (secondary tasks) have been frequently investigated in behavioral studies leading to a deterioration of working memory performance. Yet, the underlying attentional control processes are not sufficiently understood. A lateralized working memory task was frequently interrupted by either a high- or low-demanding arithmetic task and a subsequent retroactive cue indicated the working memory item required for later report. We examined the role of frontal theta (4-7 Hz) and posterior alpha power (8-14 Hz) as correlates for retroactive attentional switches between working memory representations. In particular, highly demanding interruptions decreased primary task performance compared to a control condition without interruption. This was also reflected in decreased frontal theta power and higher posterior alpha power after retro-cue presentation, suggesting decreased attentional control resources. Moreover, reduced alpha lateralization indicated an impaired refocusing on primary task information following the interruption. These results highlight oscillatory mechanisms required for successfully handling the detrimental effects of interruptions.

## 1 Introduction

“I worked all day without interruption“ is probably the least common phrase you would ever use, especially when a virus is forcing you to work from home. Interruptions (e.g., phone calls, messages or emails) are defined as secondary tasks that have been shown to deteriorate task performance, for example indicated by poorer accuracy and slower response times (Bailey & Konstan, 2006; Baron, 1986; Eyrolle & Cellier, 2000; Hodgetts & Jones, 2006). The reason for these negative effects can be attributed to higher cognitive demands, as interruptions require a switch of attention from a primary to a secondary task as well as the intention to resume the primary task afterwards (Altmann & Trafton, 2002; Brixey et al., 2007; Clapp, Rubens, & Gazzaley, 2010). Interruptions can thus be distinguished from typical dual tasking situations, as the secondary task has to be completed before the primary task is resumed. Consequently, they demand a reactivation of task-relevant information after the interruption phase (Monk, Trafton, & Boehm-Davis, 2008; Trafton, Altmann, & Ratwani, 2011).

The working memory system is of central importance in this regard, as it allows to flexibly re-organize the priority of stored mental representations (Baddeley, 2012; de Vries, Van Driel, Karacaoglu, & Olivers, 2018; Rösner, Arnau, Skiba, Wascher, & Schneider, 2020; Schneider, Göddertz, Haase, Hickey, & Wascher, 2019). This is ensured by top-down attentional control mechanisms that guide the orienting of attention towards task-relevant information and potentially lead to the inhibition of irrelevant information (Baddeley, 1996; Cowan et al., 2005; Griffin & Nobre, 2003). However, there is a great need to reconcile the understanding of attentional control processes on the level of working memory with current findings in the field of interruption research (for review: Couffe & Michael, 2017).

So far, much of the research on interruptions has been based on behavioral studies in experimental scenarios that closely resemble real-life situations (e.g., workplace-related scenarios). Important behavioral parameters in this regard are the *interruption lag*, i.e., the time between the notification of an interruption and the first performance regarding the interruption task, and the *resumption lag*, i.e., the time required to resume the performance on the primary task after an interruption (Altmann & Trafton, 2002; Trafton, Altmann, Brock, & Mintz, 2003). The interruption lag involves the preparatory processes prior to an interruption and its subsequent perception and interpretation (Baethge, Rigotti, & Roe, 2014). The resumption lag, on the other hand, includes the reactivation of goals and schemata of the primary task (Altmann & Trafton, 2002; Trafton et al., 2003). It should thus be based on attentional control processes on the level of working memory. Previous research has shown that particularly complex interruptions (e.g. math tasks; Cades et al., 2008) and interruptions in the middle of a primary task (Bailey & Konstan, 2006) led to longer resumption lags, probably because they required more cognitive resources and thereby hampered the ongoing working memory storage of mental representations related to the primary task (Cades et al., 2008; Hodgetts & Jones, 2006; Monk, Boehm-Davis, & Trafton, 2004). Recent neuroimaging studies support this notion as they reported that working memory content was not represented in neuronal activity of prefrontal cortex (PFC) regions while performing an interruption or secondary task, but had to be reactivated afterwards (Clapp et al., 2010; Mishra, Zanto, Nilakantan, & Gazzaley, 2013; Postle, Druzgal, & D’Esposito, 2003; Solesio-Jofre et al., 2012). Based on these results, the assumption was made that interruptions distort the information stored in working memory by occupying the focus of attention and thereby disrupting rehearsal processes (Bae & Luck, 2018). However, it has not yet been fully addressed how exactly attentional control processes contribute to the handling of interruptions. Particularly, the neurocognitive mechanisms involved in the switching of the focus of attention between a primary and a secondary task during the interruption and resumption lag phases have not yet been fully clarified.

To get a better idea of these mechanisms, we used a working memory paradigm with two types of interrupting arithmetic tasks (low vs. high cognitive load) and measured neural oscillations by means of the electroencephalogram (EEG) as correlates of attentional control processes. More specifically, we aimed to identify electrophysiological correlates of the preparation for the interruption (during the interruption lag) as well as the reorienting of attention to the primary task after an interruption (during the resumption lag). In this regard, two main oscillatory EEG parameters come into play: Posterior alpha (8-14 Hz) and frontal theta (4-7 Hz) oscillations. Both appear to reflect important control mechanisms required for switching the focus of attention between working memory representations (Riddle, Scimeca, Cellier, Dhanani, & D’Esposito, 2020; for review: de Vries, Slagter, & Olivers, 2020). The oscillatory power in the alpha frequency range has been shown to decrease in cortical areas linked to the processing of task-relevant stimuli and to increase in those areas representing task-irrelevant information, for example when preparing for the presentation of sensory input at a cued location (e.g., Bonnefond & Jensen, 2013; Klimesch, 2012; Roux & Uhlhaas, 2014; Thut et al., 2006; Worden et al., 2000), but also when directing the focus of attention on the level of working memory representations (e.g., de Vries et al., 2018; Hakim et al., 2019; Myers et al., 2015; Schneider et al., 2016, 2019). Also, frontal theta oscillations (4-7 Hz) have been linked to attentional control processes. They have been associated with a higher need for cognitive control (Cavanagh & Frank, 2014; Cavanagh & Shackman, 2015; Cavanagh, Zambrano-Vazquez, & Allen, 2012) and have been shown to increase when switching the focus of attention between the working memory representations of two sequential tasks (de Vries et al., 2018). In the context of interruptions, frontal theta oscillations have furthermore been studied in the interruption lag time window. For instance, Arnau and colleagues (2019) reported increased frontal theta power after a cue had given notice of the upcoming interrupting math task. Both parameters, alpha and theta oscillations, should thus be considered when investigating attentional control processes required for switching the focus of attention between the processing of the primary task and the interruption task as well as vice versa.

In the present study, the primary task required the storage of the orientation of two laterally presented bars. This process was interrupted by either of the two non-lateralized tasks (high-demanding arithmetic task vs. low-demanding number comparison task; see figure 1A). These conditions (high- and low-demanding interruption) were compared to two conditions without an interruption (prolonged fixation and early probe condition) that differed in terms of the length of the working memory storage interval. A pre-cue presented after the memory array indicated which condition would occur. After the interruption (or the intervals without interruption), a retro-cue indicated the position of the bar whose orientation had to be reported at the end of the trial.

**Figure 1.**
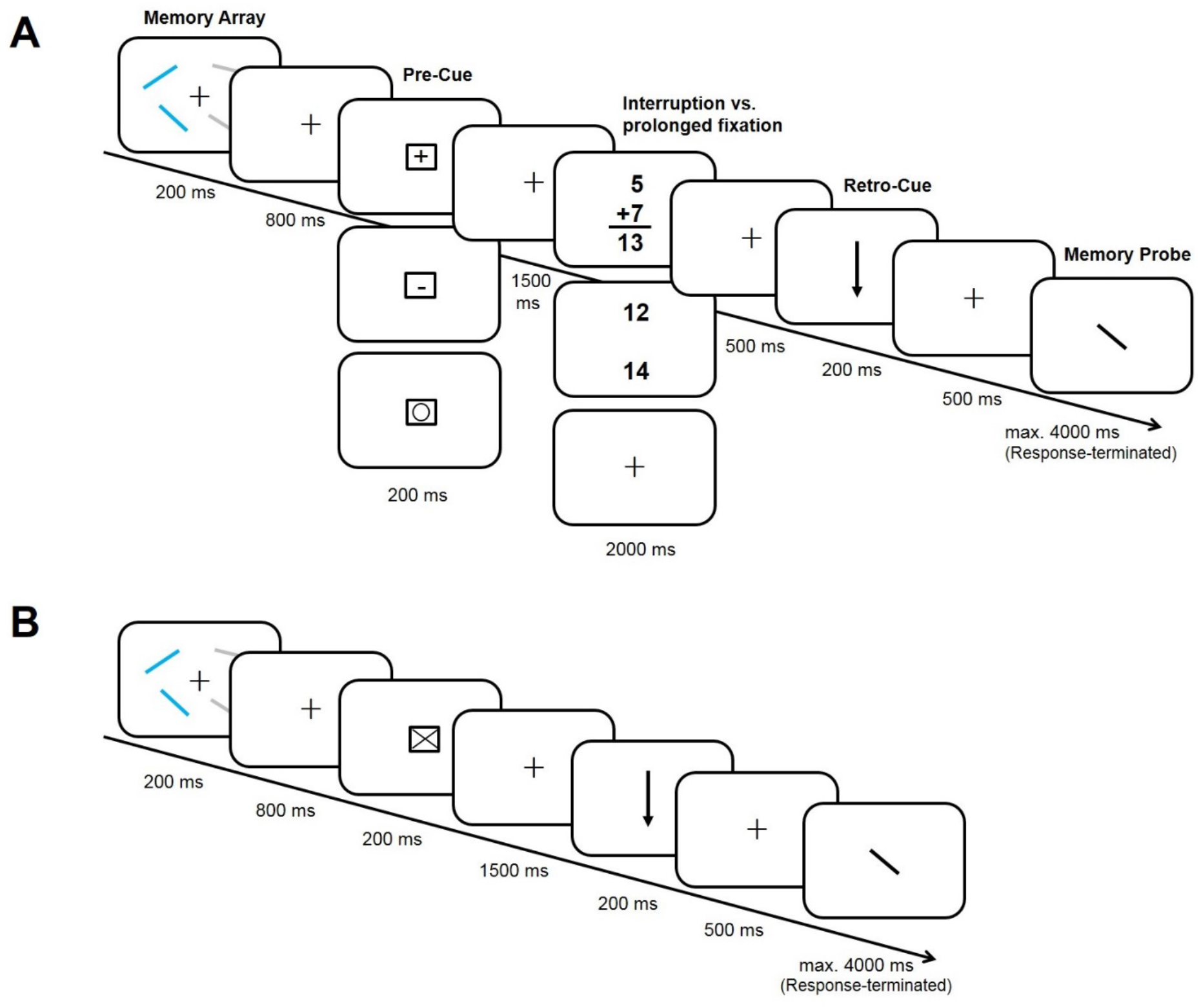
Experimental design. Each trial began with the presentation of a memory array comprising two laterally presented and randomly oriented blue bars and two irrelevant grey bars. Participants’ primary task was to remember the blue bars’ orientation of which one bar had to be reported at the end of the trial (as indicated by a retro-cue). In-between this primary task, a pre-cue indicated if and what kind of interruption would occur. Panel A depicts a trial sequence with a high-demanding arithmetic task (decide whether the math equation is correct or false; high-demanding interruption condition) a low demanding number comparison task (decide whether the lower number is larger or smaller than the upper one; low-demanding interruption condition) and a prolonged fixation cross. Panel B illustrates an additional control condition with a short storage interval (early probe condition).

We hypothesized that interruptions would negatively affect the selective retrieval of task-relevant information from working memory and therefore lead to inferior primary task performance. In line with previous behavioral studies, this impact should be more pronounced in the high-demanding interruption condition, as complex interruptions should require more attentional and cognitive resources and prevent the rehearsal of primary task information (Cades et al., 2008). On the electrophysiological level, we expected to observe stronger frontal theta power suggesting increased cognitive control in preparation for a high-demanding interruption (Arnau et al., 2019). Furthermore, an increase in theta power should be evident following the retro-cue when selecting primary task information after a delay with or without interruption. If this effect reflected the need for additional attentional control when selectively reactivating a mental representation of the primary task, it should be increased following interruptions compared to the condition without interruption (i.e., when attention had already been focused exclusively on primary task information during the delay period). However, if it rather reflected the available attentional resources for selecting the cued working memory item for later report, there should be a stronger increase in frontal theta power when participants did not have to reallocate attention to the primary task (i.e., in the conditions without interruption). Furthermore, prior research indicated that a cognitive manipulation of working memory content is also related to a suppression of oscillatory power in the alpha frequency range (~8-14 Hz) over posterior recording sites (Sauseng, Klimesch, Schabus, & Doppelmayr, 2005; Schneider, Barth, & Wascher, 2017). Referred to the current experiment, a more effective retroactive selection process in the condition without interruption should thus be reflected in a stronger suppression of posterior alpha oscillations.

Additionally, the processing of the lateral to-be-memorized stimuli in the primary task should bring about a lateralized alpha power response, with a stronger suppression at contralateral compared to ipsilateral posterior electrodes. The reactivation of primary task information during the resumption lag should accordingly be reflected by a return of this posterior alpha lateralization. Furthermore, the selection of the retroactively cued bar should again elicit a contralateral suppression of alpha power, because this process requires the selective access to the stored working memory representations (Myers et al., 2015; Poch, Campo, & Barnes, 2014; Schneider et al., 2016). These posterior alpha power asymmetries might be modulated by the type of interruption, with reduced effects indicating a less efficient reactivation of the primary task and a deficient retroactive focusing of attention on the cued working memory item, respectively.

Summarized, this study extends previous investigations by introducing electrophysiological correlates of attentional control processes during the interruption and resumption lag phases that have already been shown to be critical for the efficient handling of interruptions in more application-oriented experimental settings. We propose that interruptions, especially cognitively demanding interruptions, temporarily occupy the focus of attention and thus deteriorate working memory functions related to the primary task. More specifically, a withdrawal of the focus of attention should affect the selective focusing on primary task information and thereby degrade working memory performance. These hypothesized results should allow for gaining a better understanding of the attentional processes required for the pursuit of goal-directed behavior in the context of interfering tasks.

## 2 Results

### 2.1 Behavioral data

For the behavioral analyses, we compared performance in the primary task between the conditions with high-demanding interruptions, low-demanding interruptions, and the prolonged fixation condition (no interruption). In this regard, the angular error (see figure 2A) was measured as the degree difference between the orientation of the reported probe bar and the cued target bar of the memory array. All analyses were based only on trials in which the subjects had confirmed via button press that they had completed the interruption task and the adjustment of the probe bar.

**Figure 2.**
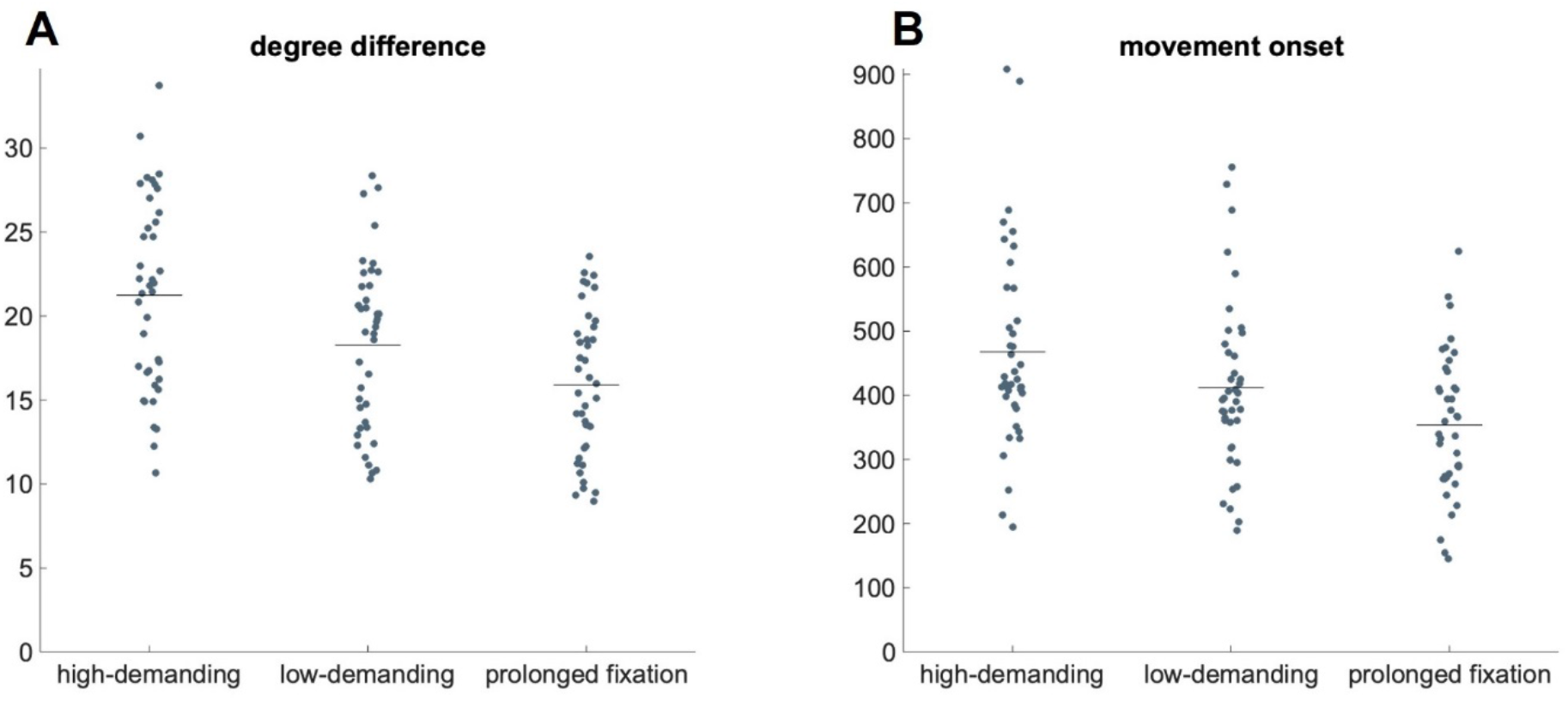
Behavioral results. 2A depicts the raw angular error separately for each condition. Each dot represents the mean angular error of one participant, while the vertical line indicates the mean of each condition. The time to mouse movement onset in ms separated for each condition is shown in 2B.

Due to a technical error (see the methods section for further details), the memory probe in the early probe condition was displayed with a 500 ms offset for 15 datasets. To ensure that the technical error did not affect any of the other three conditions we made use of an ANOVA with *Condition* (high-demanding, low-demanding, prolonged fixation) as within-subject factor and a between-subjects factor including the two samples (correct vs. erroneous early probe condition). No interactions were found for either of the behavioral parameters (angular error: *F*(2,76) = 0.15, *p* = .865, *η^2^_p_* < .01; time to mouse movement onset: *F*(2,76) = 1.03, *p* = .362, *η^2^_p_* = .03) . Thus, the following analyses are based on the data of all 40 participants (except for the comparison between the prolonged fixation and the early probe condition).

Analyses revealed significant differences regarding angular errors in the primary task between the three conditions (see figure 2A), *F*(2,78) = 66.31, *p* < .001, *η^2^_p_* = .63, ε = 0.87 (high-demanding interruption: *M* = 21.23, *SD* = 5.7; low-demanding interruption: *M* = 18.28, *SD* = 5; prolonged fixation: *M* = 15.89; *SD* = 4.29). Post-hoc *t*-tests showed a decreased accuracy in both interruption conditions compared to the prolonged fixation condition (high-demanding interruption vs. prolonged fixation: *t*(39) = 9.9, *p_adj_* < .001, *d_z_*= 1.56; low-demanding interruption vs. prolonged fixation: *t*(39) = 6.26, *p_adj_* < .001, *d_z_* =1.98), with the highest performance decrease in the high-demanding interruption condition, *t*(39) = 6.43, *p_adj_* < .001, *d_z_* = 2.03 (high-demanding interruption vs. low-demanding interruption).

Furthermore, the onset of computer mouse movement used for orientation adjustment served as a behavioral parameter for the time needed for response preparation. As shown in figure 2B, movement onset time differed between the conditions, *F*(2,78) = 36.86, *p* < .001, *η^2^_p_* = 4.86, ε = 0.49 (high-demanding interruption: *M* = 467.51 ms, *SD* = 154.27; low-demanding interruption: *M* = 411.62 ms, *SD* = 133.22; prolonged fixation: *M* = 353.73 ms; *SD* = 109.61), with faster onset times in the prolonged fixation condition compared to the high-demanding interruption, *t*(39) = −6.98, *p_adj_* < .001, *d_z_* = 2.21, and the low-demanding interruption condition, *t*(39) = −5.35, *p_adj_* < .001, *d_z_* = 1.69. Furthermore, the movement onset time was longer in the condition with high-demanding interruptions than low-demanding interruptions, *t*(39) = 4.66, *p_adj_* < .001, *d_z_* = 1.47. These result patterns indicate that behavioral performance in the primary task was not only impaired by the presence of an interruption, but also affected by the complexity of the interruption task.

The lower primary task performance following interruptions compared to the prolonged fixation condition might be related to the interruptions’ interference with working memory processes and/or to the inability to rehearse the primary task information during the delay interval. Thus, the essential question is whether a difference in primary task performance between conditions with and without interruption was due to a benefit related to the prolonged fixation condition, or to a deficit caused by the interruption. To differentiate between these possibilities, the early probe condition (i.e., cueing and probing the working memory contents without a potentially beneficial delay interval; see the methods section for further details) was included. The angular error was lower in the early probe (*M* = 14.6, *SD* = 3.91) compared to the prolonged fixation condition (*M* = 15.71, *SD* = 4.4), *t*(24) = −3.94, *p* < .001, *d_z_* = −1.58. The ability to rehearse the primary task information did therefore not benefit primary task accuracy.

Considering performance in the interruption task itself, we found higher response times to a high-demanding interruption (*M* = 1250.9 ms, *SD* = 135.4) compared to a low-demanding interruption (*M* = 885.81 ms, *SD* = 143.38), *t*(39) = 15.86, *p* < .001, *d_z_* = 2.51. There was also a lower rate of correct responses to a high-demanding (*M* = 0.82, *SD* = 0.13) compared to a low-demanding interruption (*M* = 0.97, *SD* = 0.04), *t*(39) = −7.16, *p* < .001, *d_z_* = −1.13. This indicates that the interruption condition that had been expected to be more cognitively demanding (i.e., the arithmetic task) was indeed more difficult than the number comparison task.

#### 2.1.1 Mixture Modeling

In addition to these behavioral analyses, a mixture model was fitted to the continuous performance (orientation) responses in order to analyze the precision of working memory report, as well as the probability to report the correct item (vs. reporting the wrong item or reporting on chance level; see Bays et al., 2009 and the methods section for more details). The precision of the reported orientation (kappa) differed between the three conditions, *F*(2,78) = 5.08, *p* = .008, *η^2^_p_* = .11 (see figure 3A). Post-hoc *t*-tests revealed a lower kappa for the high-demanding interruption, *t*(39) = −3.32, *p_adj_* = .002, *d_z_* = −0.53, but not for the low-demanding interruption, *t*(39) = −1.77, *p_adj_* = .084, *d_z_* = −0.28, compared to the prolonged fixation condition. The two interruption conditions did not differ from each other, *t*(39) = 1.29, *p_adj_* = .202, *d_z_* = 0.21.

**Figure 3.**
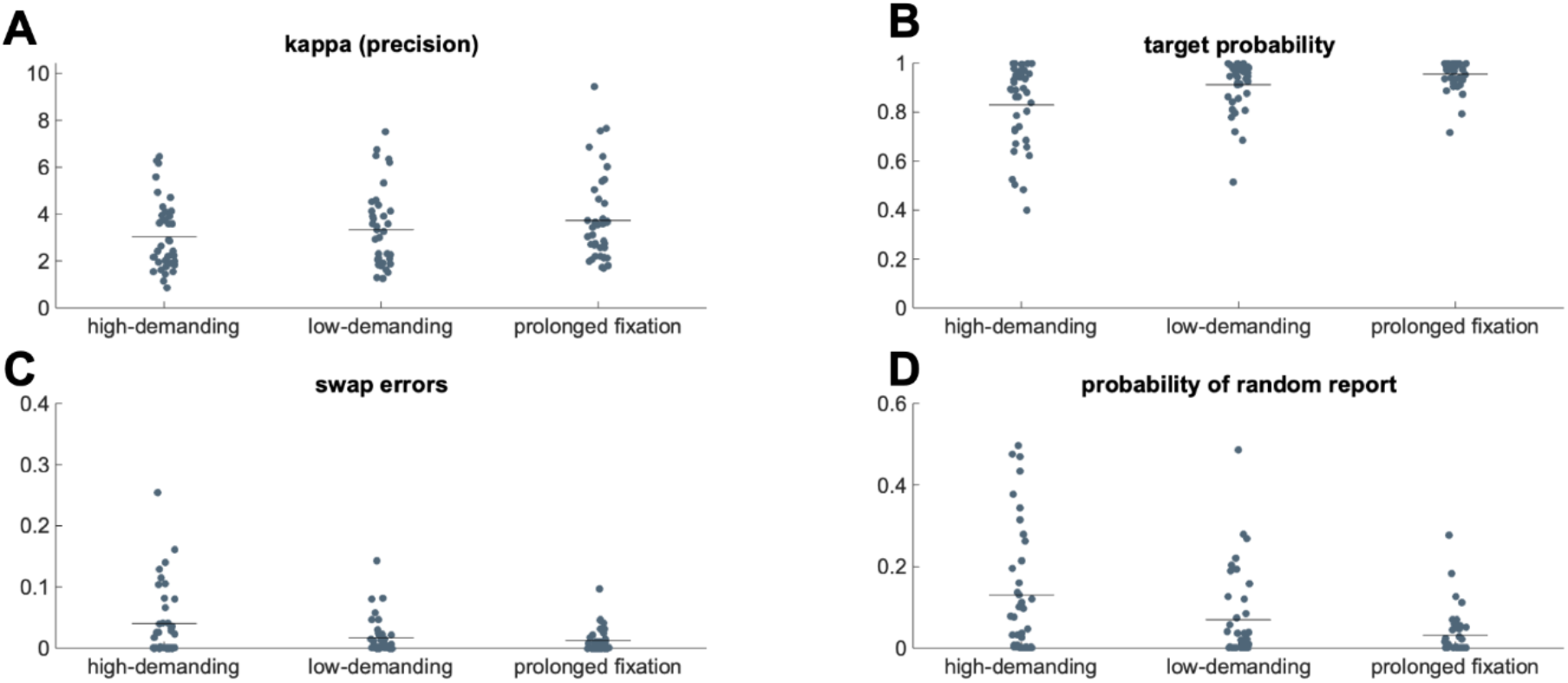
Mixture Model Parameters. In every subplot, each dot represents the mean of each participant for each condition and the vertical lines highlight the means of each condition. The precision of the mixture model parameters for each condition is shown in panel 3A. 3B depicts the mean target probability. In 3C, the probability of reporting the wrong item (swap error) is shown. The other possible source of errors is the guessing rate (probability of random report), which is depicted in panel 3D.

Furthermore, the probability of reporting the orientation of the target bar differed between the three conditions, *F*(2,78) = 24.16, *p* < .001, *η^2^_p_* = .38, ε = 0.76, (see figure 3B). In particular, the precision of target recall was reduced by a preceding interruption task (high-demanding interruption vs. prolonged fixation: *t*(39) = −5.51, *p_adj_* < .001, *d_z_* = −0.87; low-demanding interruption vs. prolonged fixation: *t*(39) = - 2.89, *p_adj_* = .006, *d_z_* = −0.45). There was also a weaker probability of target recall following a high-demanding interruption compared to a low-demanding interruption, *t*(39) = −4.87, *p_adj_* = .005, *d_z_* = −0.76. Also, the probability to report the wrong item stored in working memory (i.e., a swap error) differed between the three conditions, *F*(2,78) = 8.19, *p* < .001, *η^2^_p_* = .17, ε = 0.76. There was an increase in swap errors following high-demanding interruptions compared to low-demanding interruptions, *t*(39) = 3.12, *p_adj_* = .005, *d_z_* = 0.49, and compared to the prolonged fixation condition, *t*(39) = 3.11, *p_adj_* = .005, *d_z_* = 0.49. There was no difference in the amount of swap errors between the low-demanding interruption and the prolonged fixation condition, *t*(39) = 0.92, *p_adj_* = .362, *d_z_* = 0.14 (see figure 3C).

The lower probability to report the target item following interruptions was also based on an effect of the probability of random orientation report, *F*(2,78) = 11.66, *p* < .001, *η^2^_p_* = .23, ε = 0.87, (see figure 3D). Random report probability was higher in the high-demanding interruption condition compared to the low-demanding condition, *t*(39) = 3.16, *p_adj_* = .006, *d_z_* = 0.47. There was also a reliably higher rate of random reports in low-demanding interruption trials compared to trials without interruption, *t*(39) = 2.07, *p_adj_* = .04, *d_z_* = 0.35. Overall, these mixture modeling results point to a deterioration of reporting working memory content following interruptions that was based on both a reduced precision of the stored mental representations and a reduced probability to report the cued item, particularly after highly cognitively demanding interruption tasks.

### 2.2 EEG data

#### 2.2.1 Lateralized posterior alpha power

Figure 4 illustrates the hemispheric lateralization of posterior alpha power, which served as a correlate of attentional switches within working memory (de Vries et al., 2018; Myers et al., 2015). After processing the interruption task, a contralateral suppression of posterior alpha power that had been already observed after memory array presentation appeared (see figure 4 and table 1, time window t1). Post-hoc *t*-tests indicated that hemispheric alpha power asymmetries in both interruption conditions differed from the prolonged fixation condition (high-demanding interruption vs. prolonged fixation: *t*(39) = - 2.45, *p_adj_* = .038, *d_z_* = - 0.39; low-demanding interruption vs. prolonged fixation: *t*(39) = −2.32, *p_adj_* = .038, *d_z_* = − 0.37), but not from each other, *t*(39) = 0.19, *p_adj_* = .853, *d_z_* = −0.03. This demonstrates that participants oriented their attention back to the primary task already before retro-cue presentation.

**Figure 4.**
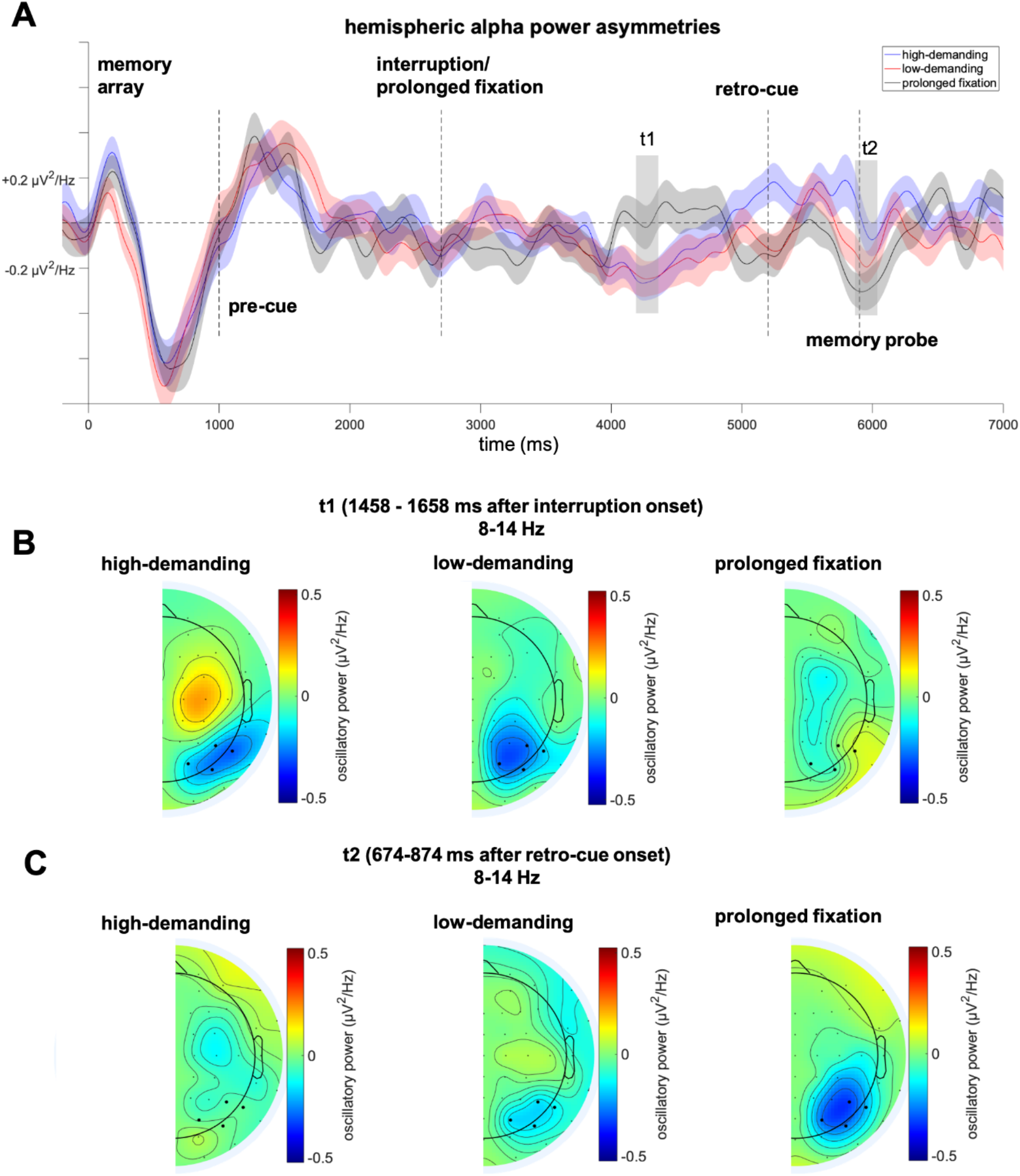
Hemispheric alpha power asymmetries. The difference wave (contra-ipsilateral to target position in the memory array) is illustrated separately for each condition (4A). The vertical lines indicate events in the trial (0 ms memory array onset, 1000 ms pre-cue onset, 2700 ms interruption onset, 5200 ms retro-cue onset, 5900 ms memory probe onset). The grey areas highlight the analyzed 200 ms time windows surrounding peaks in the grand average after each event (t1: 1458 – 1658 ms after interruption onset, t2: 693 – 893 ms after retro-cue onset). The topographical distribution for each condition of the effect in t1 is shown in panel 4B and the one for t2 in panel 4C. As the respective time windows illustrate the contralateral minus ipsilateral difference independent of hemisphere, the topographies display only half of the scalp. The analyses were based on the highlighted electrodes (PO7/PO8, P7/P8, PO3/PO4, P5/P6).

**Table 1.** Results for the non-lateralized and lateralized oscillatory power including the factor *Condition* (high-, low-demanding interruption, prolonged fixation). The time windows (t1, t2, and for the analysis of theta power also t3) provided below refer to the analyzed intervals after pre-cue, interruption, and retro-cue onset and are further depicted in figures 4 - 6.

| | df | F | $p_{adj}$ | $\eta_p^2$ | |
| --- | --- | --- | --- | --- | --- |
| **Lateralized alpha power** |  |  |  |  |  |
| **Interruption**<br>**t1: 1458 - 1658 ms** | 2,78 | 4.3 | **.034** | .1 | Condition |
| **Retro-cue**<br>**t2: 693 - 893 ms** | 2,78 | 3.4 | **.039** | .08 | Condition |
| **Non-lateralized alpha power** |  |  |  |  |  |
| **Interruption**<br>**t1: 627 - 827 ms** | 2,78 | 62.73 | **<.001** | .62 | Condition |
| **Retro-cue**<br>**t2: 342 - 542 ms** | 2,78 | 11.39 | **<.001** | .23 | Condition |
| **Non-lateralized theta power** |  |  |  |  |  |
| **Pre-cue**<br>**t1: 94 - 294 ms** | 2,78 | 1.15 | .32 | .03 | Condition |
| **Interruption**<br>**t2: 129 - 329 ms** | 2,78 | 73.91 | **<.001** | .65 | Condition |
| **Retro-cue**<br>**t3: 213 - 413 ms** | 2,78 | 27.06 | **<.001** | .41 | Condition |

Following the retro-cue, hemispheric alpha power asymmetries were evident again (see figure 4 and table 1, time window t2). Importantly, they differed significantly between the three conditions. Alpha lateralization following a high-demanding interruption, *t*(39) = 2.61, *p_adj_* = .039, *d_z_* = 0.41, but not following a low-demanding interruption, *t*(39) = 1.41, *p_adj_* = .236, *d_z_* = 0.22, was weaker than in the prolonged fixation condition. There was no significant difference between the high- and low-demanding interruption conditions, *t*(39) = 1.2, *p_adj_* = .237, *d_z_* = 0.19.

#### 2.2.2 Non-lateralized oscillatory power

Figure 5 depicts the power of alpha oscillations at posterior channels. This oscillatory response started to differ as a function of interruption condition only after interruption presentation. During interruption processing, the posterior alpha power decrease differed between the three conditions (see figure 5 and table 1, time window t1). It is trivial to note that both interruption conditions differed from the prolonged fixation condition (high-demanding interruption vs. prolonged fixation: *t*(39)= −9.12, *p_adj_* < .001, *d_z_*= −1.44; low-demanding interruption vs. prolonged fixation: *t*(39) = −7.83, *p_adj_* < .001, *d_z_* = −1.24). As can be seen in figure 5A, the difference between the high-demanding and the low-demanding interruption conditions (see time window t1), was due to a difference in latency of the alpha suppression *t*(39) = −2.41, *p_adj_* = .021 *d_z_* = −0.38. When accordingly adjusting the latencies for measuring mean alpha power (758 – 808 ms after high-demanding interruption onset; 591 – 641 ms after low-demanding interruption onset), no significant differences were found between the high-demanding and low-demanding conditions, *t*(39) = 0.41, *p* = .683, *d_z_* = 0.06.

**Figure 5.**
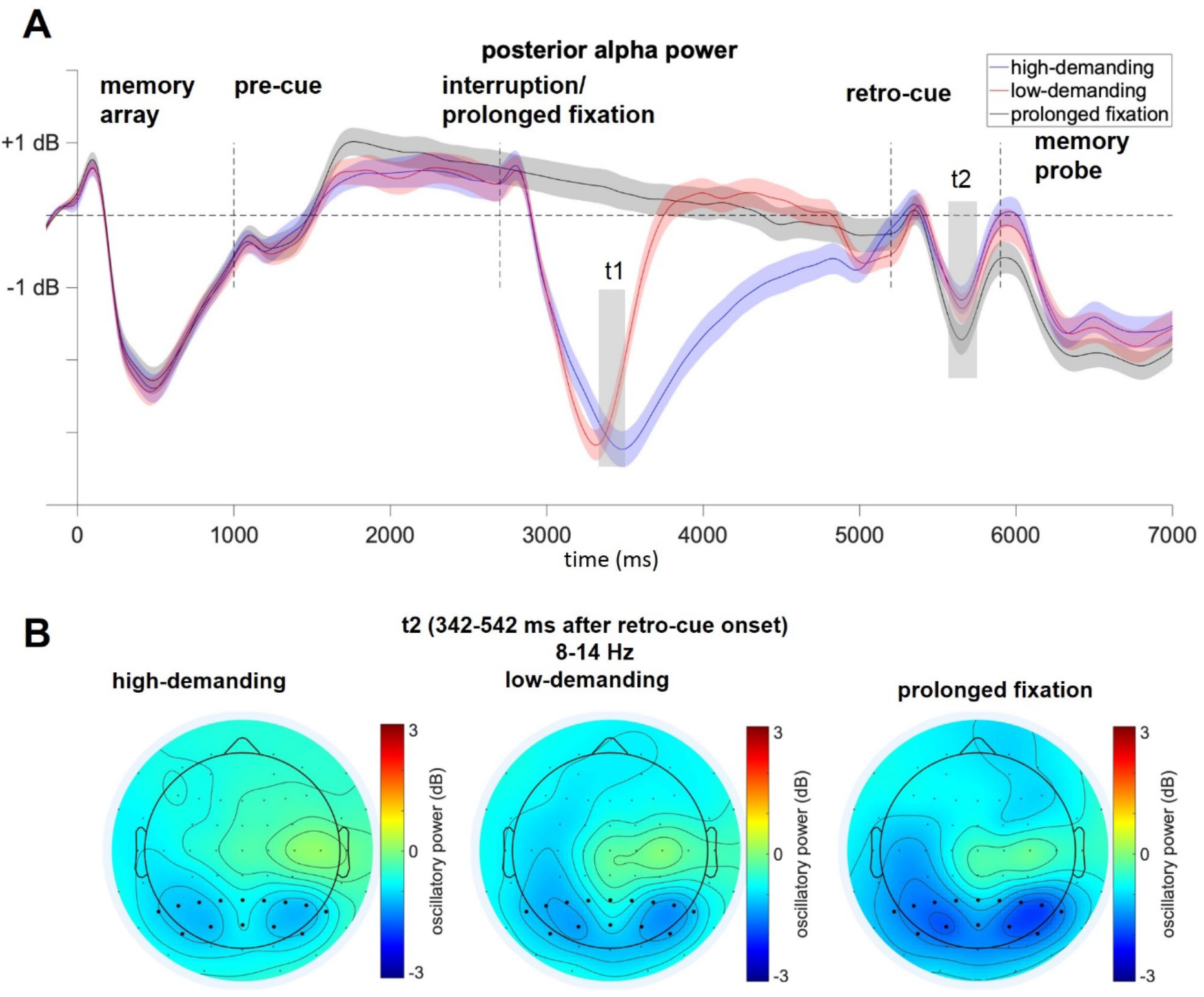
Posterior alpha power. Depicted are the ERSPs in the alpha frequency range assessed from a cluster of posterior channels separately for each condition over the whole trial (5A). The vertical lines indicate events in the trial (0 ms memory array onset, 1000 ms pre-cue onset, 2700 ms interruption onset, 5200 ms retro-cue onset, 5900 ms memory probe onset). It was tested for differences between the conditions during 200 ms time windows surrounding troughs in the grand average after each event. The respective time windows (t1: 627 – 827 ms, t2: 342 – 542 ms) are highlighted within the grey areas. The topographical distribution of relative alpha power of each condition in t1 is depicted in panel 5B, while the same is depicted in panel 5C for the time window t2.

Also, the suppression of posterior alpha power following the retro-cue differed between the three conditions (see figure 5 and table 1, time window t2). Post-hoc *t*-tests showed that alpha power decreased more in the prolonged fixation condition compared to the high-demanding, *t*(39) = 3.39, *p_adj_* < .001, *d_z_* = 0.54, and the low-demanding interruption condition, *t*(39) = 4.24, *p_adj_* < .001, *d_z_* = 1.27. However, both interruption conditions did not differ from each other, *t*(39) = 0.53, *p_adj_* = .597, *d_z_* = 0.03, indicating that this non-lateralized suppression of alpha power was sensitive to a preceding interruption, but not to the differences in its cognitive demands.

Regarding mid-frontal theta power, our results did not reveal reliable differences between the experimental conditions prior to interruption presentation (see figure 6 and table 1, time windows t1). Thus, information about the type of interruption did not elicit distinct cognitive processes to prepare for the upcoming interruption task.

**Figure 6.**
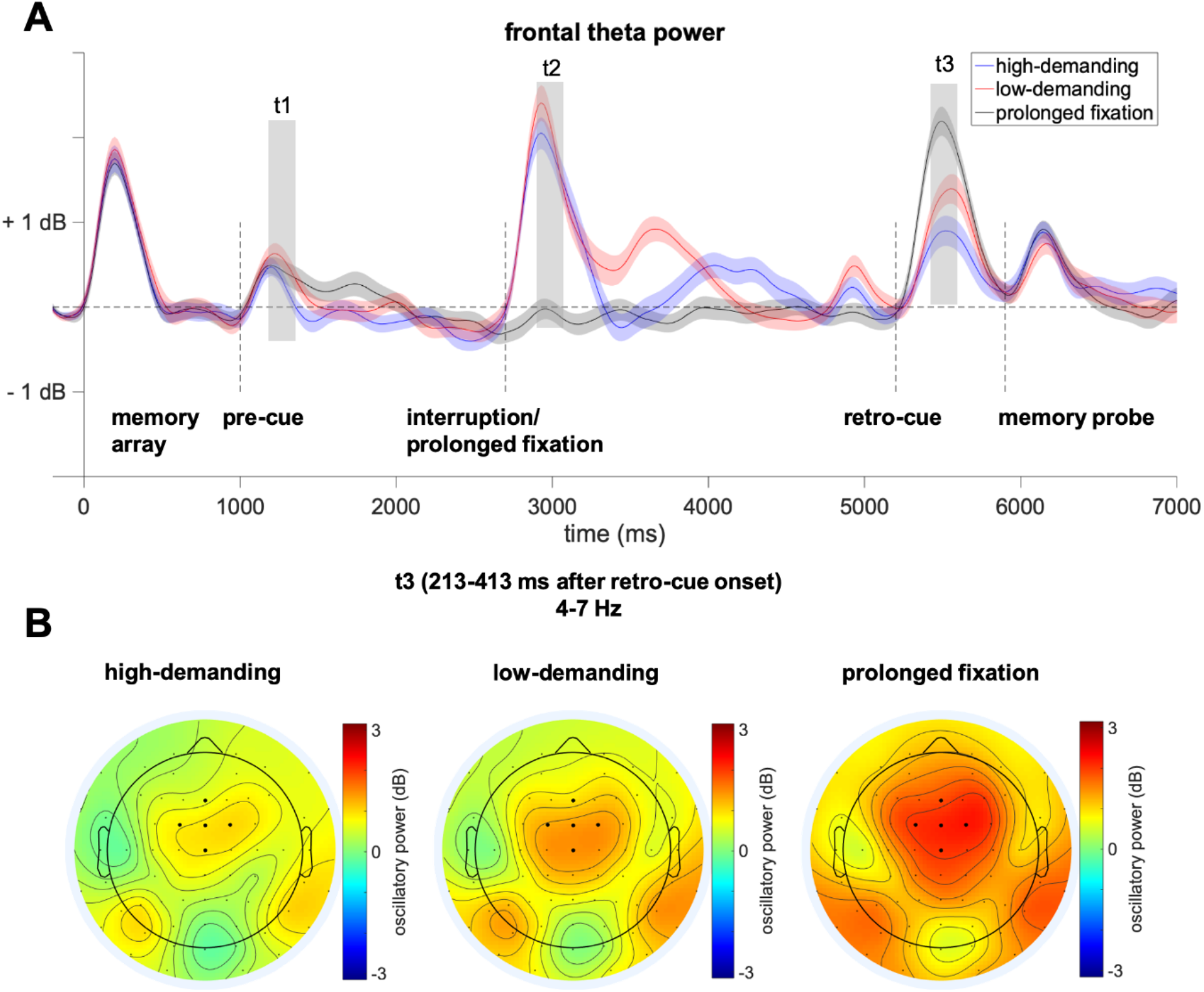
Frontal theta power. 6A displays the ERSPs at mid-frontal electrodes averaged over the theta frequency range separately for each condition. The vertical lines indicate the events within a trial (0 ms memory array onset, 1000 ms pre-cue onset, 2700 ms interruption onset, 5200 ms retro-cue onset, 5900 ms memory probe onset). It was tested for differences between the conditions in 200 ms time windows surrounding peaks in the grand average after each event. These analyzed time windows are displayed within the grey areas (t1: 94 - 294 ms, t2: 129 - 329 ms, t3: 213 - 413 ms). The topographical distribution of relative theta power for each condition in t3 is depicted in B.

Interruption onset was followed by a theta power increase (see figure 6 and table 1, time window t2). However, post-hoc *t*-tests revealed that both interruption conditions did not differ from each other in this regard, *t*(39) = −1.7, *p_adj_* = .097, *d_z_* = −054. When the retro-cue required selecting one bar from the memory array, theta power increased again. There was a main effect of *Condition* after retro-cue onset (see figure 6 and table 1, time window t3). Interestingly, paired-samples *t*-tests revealed that the theta power increase was diminished by a preceding interruption: in both, the high-demanding, *t*(39) = −6.23, *p_adj_* < .001, *d_z_* = 1.97, and the low-demanding interruption condition, *t*(39) = −4.88, *p_adj_* < .001, *d_z_*= −1.54, theta power was lower compared to the prolonged fixation condition. Furthermore, a high-demanding interruption led to a reduced theta power response following retro-cue presentation compared to the low demanding interruption, *t*(39) = −3.1, *p_adj_* = .004, *d_z_* = −0.98, showing that the extent to which a preceding interruption depleted the cognitive control resources engaged for the selection of working memory content depends on the interruptions’ complexity.

#### 2.2.3 EEG-behavior relations

We conducted further analyses to investigate to what extent the effects of oscillatory power in the alpha and theta frequency range were related to performance in the primary task. Trials were separated into high- and low-performance trials and compared to each other within each condition (see the methods section for more details). Since we had no clear hypothesis concerning the time window of an EEG-behavior relation, posterior alpha power and mid-frontal theta power was compared between high-and low-performance trials for each ERSP data point and a cluster-based permutation procedure was used to correct for multiple comparisons. The permutation test revealed a significant cluster in posterior alpha power from 1373 to 1853 ms after interruption onset in the low-demanding condition (see figure 7B). An ANOVA within this time window did not indicte a significant interaction between *Condition* and *Performance, F*(2,78) = 0.3, *p* = .741, 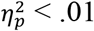. However, the effect between high- und low-performance trials after a low-demanding interruption was solely on right parietal electrodes (see figure 7D). Consequently, another ANOVA was done only considering data from four right parietal electrodes (P2, P4, P6, P8) and revealed a significant interaction between *Condition* and *Performance, F*(2,78) = 3.85, *p* = .025, 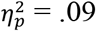. Post-hoc t-test showed that posterior alpha power was increased in high-performance compared to low-performance trials, *t*(39) = 3.06, *p_adj_* = .004, *d_z_* = 0.48. As shown in figure 7D, the maximum of this effect was evident over right parietal cortex. This relates higher alpha power after completing the interruption to a more efficient attentional refocusing on the primary task. However, there was no such difference in the high-demanding interruption condition.

**Figure 7.**
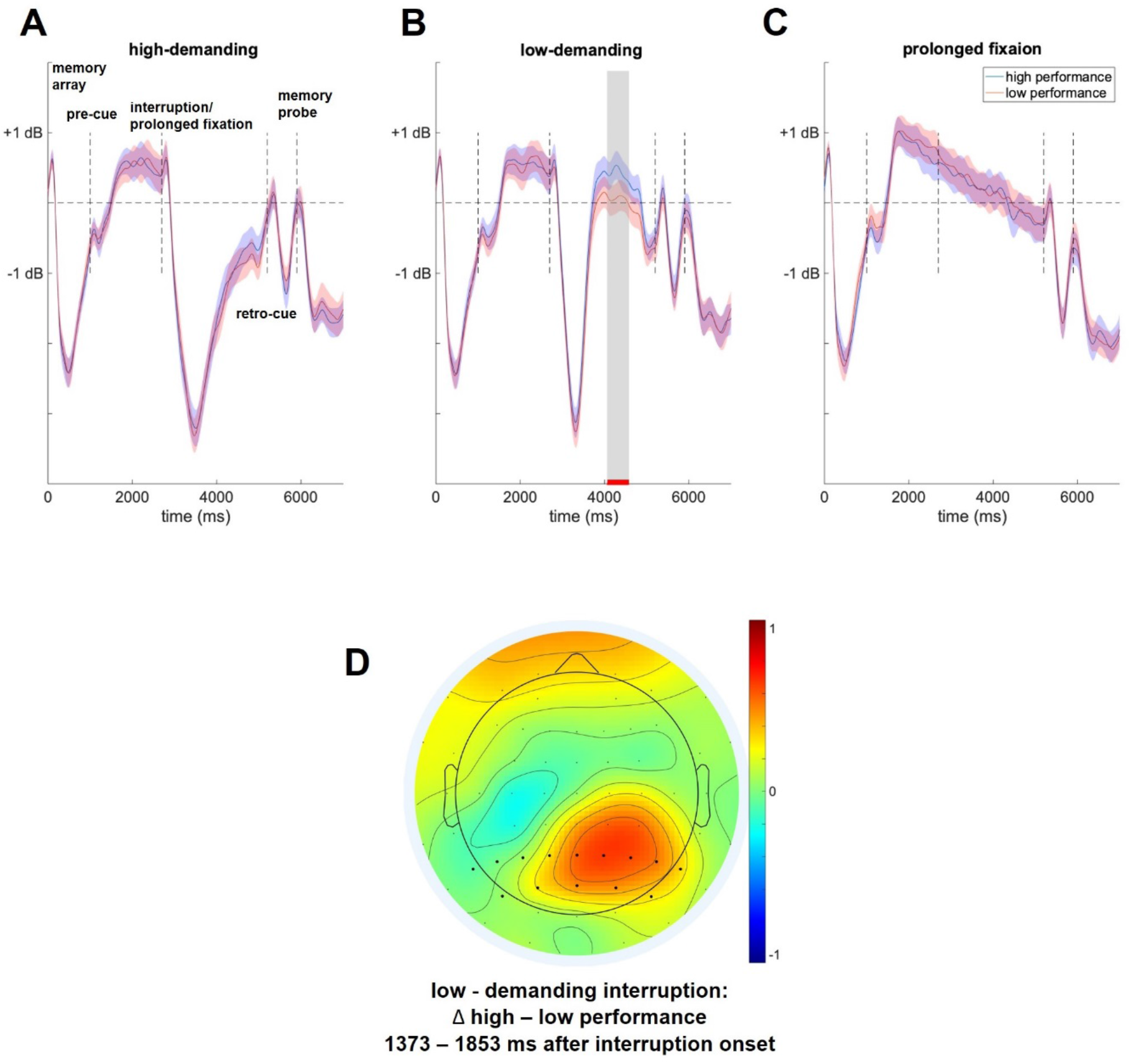
EEG-Performance relation. ERSPs are averaged over the alpha frequency range separately for high and low performance trials within each condition. The time window highlighted by the grey rectangle in the low-demanding interruption condition reached significance in the cluster-based permutation procedure. 7D shows the topographical distribution of the difference between high performance and low performance trials in the critical time window within low-demanding interruption condition.

The same analysis was run for mid-frontal theta oscillations. The cluster-based permutation procedure did not reveal any differences between high-performance and low-performance trials in any of the three conditions.

## 3 Discussion

The present study investigated how attentional control mechanisms are engaged to prevent the negative effects of interrupting tasks on working memory performance. Participants had to store the orientation of two laterally presented bars of which one had to be retrieved at the end of the trial, as indicated by a retro-cue. Crucially, prior to retro-cue presentation, this task was interrupted by a cued high- or low-demanding arithmetic task. Our results indicated that particularly high-demanding interruptions deteriorated working memory performance in the primary task, compared to conditions without interruption.

On the behavioral level, an interruption led to increased angular errors as well as to higher response onset times in the primary task compared to trials without interruption (see figure 2). In line with previous behavioral studies, this negative impact was more pronounced in the high-demanding compared to the low-demanding interruption condition (see also Cades et al., 2008; Gillie & Broadbent, 1989; Hodgetts & Jones, 2006). Importantly, the mixture model applied to the distribution of angular errors revealed in what particular way interruptions affected primary task performance. They had a negative impact on the precision of the reported working memory item and increased the probability of reporting the wrong (non-cued) working memory item (i.e., swap errors) and of random report (both probabilities contributed to a lower probability to report the cued bar’s orientation). Again, the most detrimental influence was evident following the high-demanding interruptions. This is in line with a study by Makovski and Pertzov (2015) which reported interruptions to particularly increase swap errors and impair report precision. It is therefore reasonable to assume that interruptions affect both the precision of a working memory item (i.e., the content level) and the ability to retrieve working memory content based on contextual information. The latter effect might be due to either a complete loss of primary task information (leading to a random orientation adjustment) or to a weakened binding of working memory content to (spatial) context information (leading to a higher rate of swap errors; see also Schneegans & Bays, 2019).

To get a better understanding of the attentional control functions involved in the efficient handling of interruption tasks, we further focused on frontal theta (4-7 Hz) and posterior alpha oscillations (8-14 Hz), as both oscillatory parameters have been related to the re-allocation of the focus of attention in working memory (e.g., de Vries et al., 2018; Riddle et al., 2020; Rösner et al., 2020; Schneider et al., 2019). The suppression of alpha power contralateral to the relevant side of the memory array indicated that the participants oriented their attention towards the relevant blue bars. This contralateral alpha power suppression returned after performing the interruption task, suggesting that participants shifted the focus of attention back to the primary task as soon as they completed the interruption task. This finding highlights alpha power lateralization as a reliable indicator for the resumption of the primary task and thus constitutes an electrophysiological correlate of the resumption lag that should be more precise than the previous approaches based on behavioral parameters (Trafton et al., 2003). In the current experiment, this lateralized effect, however, did not differ as a function of the difficulty of the interruption task (see figure 4). This could indicate that, with regard to the reactivation of the primary task in working memory, participants were guided more by the time requirements of the interruption task than by the individual processing time required for the interruption. In other words, this could mean that, at least in the highly demanding interruption task, a reactivation of the primary task information took place to some extent already during the interruption phase.

The subsequent retro-cue presentation was again followed by a contralateral suppression of alpha power that can be linked to the attentional selection of the cued memory array item (de Vries et al., 2018; Schneider, Barth, Getzmann, & Wascher, 2017; Schneider et al., 2019). While we found no significant difference between high- and low-demanding interruption trials, the reduced alpha lateralization compared to the prolonged fixation condition still suggests that a withdrawal of the focus of attention by means of an interruption impaired the ability to subsequently select the cued working memory item. This is also in line with the current behavioral results and earlier investigations showing a deterioration of the probability to retrieve the correct working memory item after interruptions (see also Makovski & Pertzov, 2015).

While the posterior alpha power lateralization can thus be used as a marker for the efficiency of retroactive attentional selection of primary task information after an interruption, changes in non-lateralized oscillatory power during a working memory task are rather associated with the more general availability of attentional control resources (Cavanagh & Frank, 2014; Cavanagh & Shackman, 2015; Cavanagh et al., 2012). With respect to retro-cue presentation, our results revealed significantly lower frontal theta power after interruptions compared to the conditions without a preceding interruption, with the lowest theta power following high-demanding interruptions (see figure 6). This result pattern is consistent with our hypothesis that frontal theta power in the context of retroactive attentional orienting reflects the ability to exert attentional control on the level of working memory rather than the actual need for such a process (cf. Cavanagh et al., 2012; Cavanagh & Frank, 2014; Zanto et al., 2010). Similar to previous findings showing decreased theta power for mental representations that are difficult to retrieve from memory (Klimesch et al., 2006; Spitzer, Hanslmayr, Opitz, Mecklinger, & Bäuml, 2009), our results suggest that in particular cognitively more demanding interruptions led to reduced cognitive control for selecting the cued working memory item. Importantly, this decrease in mid-frontal theta power after retro-cue presentation in the interruption conditions went along with a reduced posterior alpha power suppression. This is in line with previous research showing that both correlates play a crucial role in prioritizing task-relevant information and potentially in suppressing information that is no longer relevant in order to successfully guide behavior (de Vries et al., 2018; Nelissen, Stokes, Nobre, & Rushworth, 2013; Riddle et al., 2020).

We furthermore identified mechanisms relevant for the efficient handling of the detrimental effects of interruptions. Increased alpha power over the right parietal cortex following the response to the low-demanding interruption task was associated with high accuracy in the primary task. In accordance with previous studies suggesting a relation between alpha oscillations and the inhibition of irrelevant information (e.g. Bonnefond & Jensen, 2013; Händel et al., 2011; Rösner et al., 2020; Wöstmann et al., 2019), this alpha power increase might reflect the removal of mental representations of the interrupting arithmetic task from working memory after interruption processing had been completed (in the sense of an entire completion/closure of processing the interruption task). Importantly, in the high-demanding condition, alpha power in the resumption phase did not differ as a function of primary task performance and did not return to baseline level prior to retro-cue presentation (see figure 7). We therefore assume that the processing of the highly demanding arithmetic task took too much time to ensure a chronologically well-ordered completion of the interruption and resumption of the primary task. These findings are in line with a recent investigation by Wang, Theeuwes, & Olivers (2018), showing that more time to prepare for a memory test after an interruption benefits performance. In a set of experiments, it was shown that memory probe presentation after an interrupting task interfered with the memorized item when both stimuli were similar (i.e., memory item and probe presented as randomly oriented gratings) and that this effect could be compensated by providing additional processing time between the interruption task and the presentation of the memory probe. An essential recommendation for the handling of interruptions can be given based on these results and the current electrophysiological findings: The time available for resuming the primary task is a critical factor for counteracting the detrimental effects of interruptions on working memory performance. Additional time might allow for a complete inhibition of mental representations of the interruption and allow for a better attentional preparation for resuming the primary task, thereby contributing to a better performance.

In order to study the preparatory attentional processes for coping with the impact of interruptions, the present study made use of pre-cues prior to the interruption tasks. This has been shown to enable the activation of interruption schemata, thereby facilitating the resumption of the primary task (Brudzinski, Ratwani, & Gregory Trafton, 2007; Ratwani, Andrews, McCurry, Trafton, & Peterson, 2007). Hence, stronger theta power reflecting increased cognitive control in preparation for particularly high-demanding interruptions was expected (see Arnau et al., 2019). However, in the present study, we did not find significant differences in oscillatory correlates of preparatory attentional processes between the three conditions. One possible explanation for this lack of an effect could be that there was not enough time to engage in preparatory cognitive control processes based on the information provided by the pre-cues (see also Andrews et al., 2014). Future research should make use of different types of pre-cues and different stimulus-onset asynchronies (SOAs) in order to allow for investigating distinct preparatory processes during the interruption lag.

In summary, the present study demonstrated that interruptions, in particular cognitively highly demanding interruptions, impaired the resumption of a working memory task. This was indicated by both a lower precision of the retrieved working memory item and a lower probability to report the cued orientation (as a consequence of a higher swap error and random report rate). On the EEG level, lower frontal theta power and higher posterior alpha power after retro-cue presentation indicated fewer available attentional control resources following interruptions. These oscillatory effects were followed by a reduced alpha power lateralization relative to target position, indicating an inferior capability to retroactively select task-relevant information of the primary task after an interruption. These results thus extend previous findings by providing electrophysiological evidence for the assumption that interruptions affect attentional control processes and thereby hamper the reactivation of mental representations in working memory (e.g., Bae & Luck, 2018). Our task design further allowed for using posterior alpha power asymmetry as a reliable indicator for the resumption lag independent of the time required to execute the first action in the primary task. This could turn out to be a very important parameter for future investigations, as not all tasks require immediate action when resuming after an interruption. Finally, we revealed a cognitive mechanism that might be involved in the compensation of the detrimental impact of interruptions, i.e., inhibiting irrelevant secondary task information in the resumption phase. Further investigations should have a closer look at the dependency of this mechanism on the time available before resuming the primary task. This should help to provide more precise recommendations on how to deal with interruptions, for example in the context of the design of working environments.

## 4 Methods

### 4.1 Participants

A total of 44 participants were tested (*M(age)* = 24.02, *SD* = 3.49, range = 19 - 30, 27 females). Working memory recall exceeding 30% of the maximal angular error (i.e., mean angular error exceeding 27 degrees) in the prolonged fixation condition led to the exclusion of three participants. Furthermore, one participant did not follow the task instructions (i.e., remembering the bars always on the left side) and was thus excluded. Therefore, the data of 40 participants (*M(age)* = 23.75, *SD* = 3.2, range = 19-30, 25 females) were included in the analysis. All participants were right-handed according to a handedness questionnaire (adapted from Oldfield, 1971). None of them reported any known neurological or psychiatric disease and color vision was confirmed by means of the Ishihara Test for Color Blindness. Participants were compensated with 10€ per hour or course credit and gave written informed consent prior to the beginning of the experiment. The study was in accordance with the Declaration of Helsinki and approved by the local ethics committee of the Leibniz Research Centre for Working Environment and Human Factors.

### 4.2 Apparatus, stimuli and procedure

The experiment took place in an electrically shielded, dimly-lit chamber. Participants were seated in front of a 22-inch CRT monitor (100 Hz; 1024 x 768 pixels) at a 150 cm viewing distance. Stimuli presentation was controlled by a ViSaGe MKII Stimulus Generator (Cambridge Research Systems, Rochester, UK).

Prior to the main experiment, participants completed three training sessions to get acquainted with the task. The first one trained the participants in adjusting the memory probe orientation using the computer mouse. Two randomly oriented bars (1° by 0.1° visual angle) were presented in 2.12° distance to the left and right of a central fixation cross. One bar was presented in black (RGB: 0,0,0) and had to be rotated to match the same orientation as the second grey bar (RGB: 140, 140, 140) by moving the computer mouse horizontally. This orientation adjustment had to be completed within 4000 ms. The mean angular error had to be below 18° in the last 50 trials of a block of 150 trials. All participants reached this performance level within the first practice block. In the second training session, participants performed the primary task of the main experiment without interruptions (28 trials). Afterwards, in a third training session, participants performed 36 trials in order to practice the high-demanding interruption (12 trials), the low-demanding interruption (12 trials), and the prolonged fixation condition (12 trials).

For the main experiment, a black fixation cross (1.7° x 1.7°) was presented centrally on the screen during memory array presentation and the delay intervals. Participants were instructed to fixate on the cross while it was present. Each trial began with the presentation of a memory array, which was composed of two randomly oriented blue bars (luminance: 52cd/m^2^, RGB: 0, 176, 255, size: 0.1° x 1.0°) on either the left or right side of the fixation cross (see figure 1). The minimum difference in orientation of these bars was 15°. Grey bars (luminance: 52cd/m2, RGB: 140, 155, 126, size: 0.1° x 1.0°) were presented on the other side of the fixation cross to avoid laterally imbalanced sensory input. Participants were instructed to memorize only the orientation of the two blue bars.

The memory array was followed by a cue (size of the rectangle surrounding it: 3.44° x 2.58° visual angle) which indicated if the primary task was going to be interrupted or not (hereafter referred to as pre-cue). Two different interruption types were introduced: A plus sign (size: 1.7° x 1.7°) indicated that the primary task was interrupted by a highly cognitively demanding arithmetic task (high-demanding interruption; ¼ of all trials). Participants had to indicate whether this equation was solved correctly or not (which was the case in 50% of the trials) by pressing the left or right computer mouse button (index vs. middle finger). To ensure a comparable difficulty across all trials within the same condition, the sum of the arithmetic task was always a two-digit number and the task itself consisted of two one-digit numbers. Moreover, a pre-cue presented as a minus sign (size: 1.7° x 1.7°) indicated a low demanding number comparison task (low-demanding interruption; ¼ of all trials). Two different single digit numbers, which always added up to a two-digit number, were presented as a vertical array. Participants had to indicate by mouse click (left vs. right button) whether the smaller number was presented at the top or at the bottom. Both interruption tasks (high- and low-demanding interruptions) were presented centrally for 2000 ms and responses were recorded only during this interval. The assignment of response keys to experimental conditions was counterbalanced across participants. Additionally, in a control condition, a circle (size: 1.7° x 1.7°) indicated that there was no interruption but the presentation of the fixation cross was prolonged for 2000 ms (¼ of all trials), resulting in an equal trial duration in all three conditions.

Moreover, a fourth condition without interruption was introduced (see figure 1B) to rule out the possibility that the hypothesized performance differences between the conditions with and without interruption were rather based on a benefit in the prolonged fixation condition resulting from the possibility to rehearse the working memory material than on a deficit resulting from the interruptions. In one-fourth of the trials, a pre-cue presented as an “X” (size: 3.44° x 2.58° visual angle) indicated that the primary task was to be continued at the same time the interruption tasks were presented (i.e., early probe condition). This condition was not included in the EEG analyses due to a lack of comparability regarding the different temporal structure. Please also note that due to a technical error (i.e., the retro-cue was presented 500 ms later than intended) this condition could not be analyzed for 15 out of 40 participants.

Following the interval with or without interruption, a centrally presented retro-cue (an arrow with 0.69° x 1.12° visual angle) indicated one of the blue bars to be the target. At the end of each trial, a black bar in random orientation was presented as a memory probe and had to be adjusted to match the orientation of the target bar by moving the computer mouse horizontally. Participants were instructed to indicate by button press (left button of the computer mouse) when they considered the orientation adjustment as completed. The adjusted memory probe remained presented for 200 ms after this button press had occurred. A trial was counted as ‘incomplete’, when this button press did not occur within 4000 ms after memory probe presentation. The inter-trial interval varied randomly in steps of 1 ms between 500 and 1000 ms and started either 200 ms after a button press or directly after the maximum response time window. The 480 trials were divided into six blocks, separated by breaks of around two minutes to prevent fatigue during the experiment. Including the preparation for the EEG recording, the whole experimental session took approximately 4 h.

### 4.3 Behavioral analysis

Only trials in which participants responded to the primary task as well as to the interruption task (both as indicated by button press) were considered for further analyses. All responses faster than 150 ms to either the primary or interruption task were considered as premature responses. These trials were excluded from the analysis. Parameters for behavioral performance were the difference in orientation between the adjusted memory probe and the cued bar (i.e., the angular error) and the time required for starting the orientation adjustment following memory probe presentation (i.e., computer mouse movement onset time). For both dependent variables, separate repeated-measures analyses of variance (rm-ANOVA) including the within-subject factor *Condition* (high-demanding interruption, low-demanding interruption, prolonged fixation) were conducted. These analyses were based on the data from all 40 participants, despite the technical error in the early probe condition. This can be considered acceptable because the performance in the primary task and the effect of the interruption conditions on the behavioral performance did not differ between the two samples of the early probe condition as a mixed ANOVA with the within-subject factor *Condition* (high-demanding interruption, low-demanding interruption, prolonged fixation) and a between-subject factor including the two samples (correct vs. erroneous early probe condition) revealed (see results section). The early probe condition without the technical error served as a control condition and was therefore only compared to the prolonged fixation condition within a sample of 25 participants (using a paired-samples *t*-test). Furthermore, accuracies and response times were compared between the two interruption tasks by means of paired-samples *t*-test to check whether high-demanding interruptions were indeed more cognitively demanding than low-demanding interruptions.

In order to provide a more detailed estimation of the effects of interruptions on working memory performance, we further fitted a mixture model to the angular error distributions (see Bays et al., 2009). The mixture model provided four parameters for each participant within each experimental condition (high-demanding interruption, low-demanding interruption, prolonged fixation): the probability of reporting the cued (target) item, the probability of reporting the non-cued item (swap errors), the probability of random orientation report, and the parameter *kappa* which indicates the precision of the reported working memory representation. Each model component was analyzed in a rm-ANOVA with the within-subject factor *Condition* (high-demanding interruption, low-demanding interruption, prolonged fixation).

### 4.4 EEG data recording, preprocessing and analyses

The EEG was recorded with a sampling rate of 1000 Hz from 64 Ag/AgCl passive electrodes (Easycap GmbH, Herrsching, Germany) that were arranged across the scalp in an extended 10/20 scalp configuration (Pivik et al., 1993). A NeurOne Tesla AC-amplifier (Bittium Biosignals Ltd, Kuopio, Finland) was used for recording data with a 250 Hz low-pass filter. Impedances were kept below 10 kΩ. The midline electrode FCz served as reference electrode and the ground electrode was located at AFz.

MATLAB (R2019b) and the EEGLAB toolbox (Delorme & Makeig, 2004) were used for further data processing. EEG data were filtered with a 1 Hz high-pass filter and a 30 Hz low-pass filter before channels with kurtosis exceeding 10 SD were rejected (*M* = 3.5, *SD* = 1.93) based on a channel rejection tool implemented in EEGLAB. After re-referencing the data to common average of all non-rejected channels, the data were segmented into epochs ranging from 700 ms before to 7500 ms after memory array onset. Subsequently, trials containing gross artifacts were rejected based on an automatic rejection procedure implemented in EEGLAB (threshold = 500 μV, probability threshold = 5 SD, max. % of trials rejected per iteration: 5%). Afterwards, the data were down-sampled to 250 Hz prior to an Independent Component Analysis (ICA) to reduce computation time. Independent components (ICs) related to artifacts such as eye movements, eye blinks or generic data discontinuities were identified using the ADJUST algorithm (Mognon, Jovicich, Bruzzone, & Buiatti, 2011). Subsequently, the IC weights were transferred to the original low-pass and high-pass filtered 1000 Hz data. Then, single dipoles based on a spherical head model were fitted to the ICs by applying the EEGLAB plug-in DIPFIT. ICs identified by ADJUST and those with a residual variance of the dipole solution exceeding 40% were excluded from the data. The continuous EEG data were then again segmented into epochs from 700 ms before to 7500 ms after memory array presentation. The baseline was set to the 200 ms interval preceding memory array presentation.

An automatic artifact rejection procedure run on the pruned data (threshold limit: 1000, probability threshold: 5 SD, max. % of trials rejected per iteration: 5%) led to a rejection of 80.38 trials on average (range = 42 - 129, *SD* = 23.71). Finally, the rejected channels were re-computed by means of a spherical spline of the surrounding channels. Only ‘complete trials’, meaning those trials with both a response to the interruption and the primary task within the pre-defined intervals (interruption: 150 – 2000 ms after interruption onset; primary task: 150 – 4000 ms after memory probe onset) were included in the following analyses.

#### 4.4.1 Time-frequency analyses: Lateralized posterior alpha power

Spectral power or event-related spectral perturbation (ERSP; Delorme and Makeig, 2004) was computed by convolving three-cycle complex Morlet wavelets with each epoch of the EEG data. Frequencies ranged from 4 to 30 Hz in 52 logarithmic steps. The number of cycles used for the wavelets increased half as fast as the number of cycles in the respective fast-fourier transformation (FFT). This resulted in three cycle wavelets at the lowest frequency (4 Hz) and 11.25 cycles at the highest frequency (30 Hz). Resulting epochs contained 400 time points ranging from 282 ms before to 7082 ms after memory array onset.

Hemispheric asymmetries in posterior alpha power were calculated without baseline normalization at electrodes PO7/8, PO3/4, P7/8, and P5/6 in line with similar studies focusing on attentional selection processes in working memory (e.g., Rösner et al., 2020; Schneider et al., 2019). The contralateral and ipsilateral signal was computed by averaging across leftsided electrodes when target bars were presented on the right side and right-sided electrodes when targets were presented on the left side (contralateral). The same was done for left-sided electrodes when targets were presented on the left side and right-sided electrodes when targets were presented on the right side (ipsilateral). The contralateral minus ipsilateral difference in oscillatory power (μV^2^/Hz) was computed separately for each condition (high-demanding interruption, low-demanding interruption, prolonged fixation) and dataset. Troughs in the difference wave averaged across conditions were searched following pre-defined events of interest: interruption onset and retro-cue onset. Within a 200 ms time window centered on each trough, differences between conditions were identified using a rm-ANOVA including the within-subject factor *Condition* (high-demanding interruption, low-demanding interruption, prolonged fixation).

#### 4.4.2 Time-frequency analyses: Non-lateralized oscillatory power

For analyses of non-lateralized oscillatory power, the interval before memory array onset was defined as an oscillatory baseline. To assess the processing of working memory representations in posterior areas, the oscillatory power in the alpha frequency range (8 – 14 Hz) was calculated at a large electrode cluster comprising a set of parietal and parietooccipital channels (PO7, PO8, PO3, PO4, POz, P1, P2, P3, P4, P5, P6, P7, P8). Furthermore, oscillatory power in the theta frequency range (4 – 7 Hz) at a cluster of mid-frontal channels (FCz, FC1, FC2, Fz, Cz) was assessed as a correlate of higher-level cognitive control functions (Arnau et al., 2019; Cavanagh & Shackman, 2015). For statistical analyses of non-lateralized EEG data, the grand average across conditions was again computed separately for each frequency range. Time windows of 200 ms were centered on each trough for the analysis of posterior alpha power and on each peak for the analysis of frontal theta power. A rm-ANOVA was computed with the within-subject factor *Condition* (high-demanding interruption, low-demanding interruption, prolonged fixation) for each frequency range to identify differences between the three conditions.

#### 4.4.3 EEG-behavior analysis

Further, we analyzed if electrophysiological parameters assessed from a posterior electrode cluster (PO7, PO8, PO3, PO4, POz, P1, P2, P3, P4, P5, P6, P7, P8) were related to the accuracy of probe adjustment in the primary task. Using a median split, the data were separated into high-versus low-performance trials. Importantly, the median was calculated within each experimental condition (six conditions resulting from three interruption conditions and left vs. right target side) and for each participant individually. To adjust for multiple comparisons, a cluster-based permutation statistical approach was applied. Within each condition (high-demanding interruption, low-demanding interruption, prolonged fixation), the high-vs. low-performance condition labels were randomly assigned to each dataset and this was repeated 1000 times. On each iteration, a within-subject *t*-test was run for each data point resulting in a time points (400) x permutations (1000) matrix. The size of the largest cluster with *p*-values < .05 was assessed for each permutation. Performance differences in each condition were considered significant if the size of a *p*-value cluster < .05 in the original data was larger than the 95^th^ percentile of the permutation-based distribution of cluster sizes. Further ANOVAs regarding oscillatory power focused on the identified significant time windows included the within-subject factors *Condition* (high-demanding interruption, low-demanding interruption, prolonged fixation) and *Performance* (high, low). It was only tested for the interaction between both factors. One ANOVA was done on the data obtained from the whole posterior cluster and another one focused solely on data from right parietal electrodes, which revealed the highest difference between low- and high-performance trials (P2, P4, P6, P8).

### 4.5 Inferential statistics and effect sizes

For all statistical analyses regarding the behavioral and EEG data, Greenhouse-Geisser correction was applied when sphericity of the data was violated (indicated by ε). As a measure of effect size regarding the ANOVA, partial eta squared 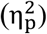 was computed. Cohens *d_z_* was used as a measure of effect size for within-subject *t*-tests. Post-hoc comparisons between the three experimental conditions required correction for multiple comparisons and therefore made use of the false discovery rate (FDR) procedure (adjusted *p*-values /*p_adj_* are reported in this regard; see Benjamini and Hochberg, 1995). Additionally, the FDR procedure was also used for correcting for multiple testing in the different time windows in the EEG analysis.

## 5 Acknowledgements

The authors would like to thank Tobias Blanke for programming the experiment, Pia Deltenre, Barbara Foschi and Katrine Bergeron for assistance in collecting the data, and Laura Klatt for proofreading the manuscript.

